# A Consistent Estimator of the Evolutionary Rate

**DOI:** 10.1101/008714

**Authors:** Krzysztof Bartoszek, Serik Sagitov

## Abstract

We consider a branching particle system where particles reproduce according to the pure birth Yule process with the birth rate λ, conditioned on the observed number of particles to be equal *n.* Particles are assumed to move independently on the real line according to the Brownian motion with the local variance σ^2^. In this paper we treat *n* particles as a sample of related species. The spatial Brownian motion of a particle describes the development of a trait value of interest (e.g. log-body-size). We propose an unbiased estimator 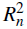 of the evolutionary rate *ρ*^2^ =σ^2^/λ. The estimator 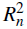 is proportional to the sample variance 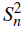 computed from *n* trait values. We find an approximate formula for the standard error of 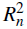 based on a neat asymptotic relation for the variance of 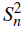.

## 1 Introduction

Biodiversity within a group of *n* related species could be quantified by comparing suitable trait values. For some key trait values like log body size, researchers apply the Brownian motion model proposed by Felsenstein [1985]. It is assumed that the current trait values 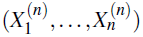 have evolved from the common ancestral state *X*_0_ as a branching Brownian motion with the local variance σ^2^. Given a phylogenetic tree describing the ancestral history of the group of species the Brownian trajectories of the trait values for sister species are assumed to evolve independently after the ancestor species splits in two daughter species. The resulting phylogenetic sample 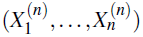 consists of identically distributed normal random variables with a dependence structure caused by the underlying phylogenetic signal.

A mathematically appealing and biologically motivated version of the phylogenetic sample model assumes that the phylogenetic tree behind the normally distributed trait values 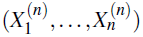 is unknown. As a natural first choice to model the unknown species tree, we use the Yule process with birth rate *λ* [see Yule, 1924]. Since the phylogenetic sample size is given, *n,* the Yule process should be conditioned on having *n* tips: such conditioned branching processes have received significant attention in recent years, due to e.g. Aldous and Popovic [2005], Gernhard [2008], Mooers et al. [2012], Stadler [2009, 2011], Stadler and Steel [2012]. This ”tree-free” approach for comparative phylogenetics was previously addressed by Sagitov and Bartoszek [2012] and Crawford and Suchard [2013], [much earlier Edwards, 1970, used a related branching Brownian process as a population genetics model].

In our work we show that a properly scaled sample variance is an unbiased and consistent estimator of the compound parameter *ρ*^2^ = σ^2^/λ which we call the evolutionary rate of the trait value in question. Our main mathematical result, Theorem 2.1, gives an asymptotical expression for the variance of the phylogenetic sample variance. This result leads to a simple asymptotic formula for the estimated standard error of our estimator. Our result is in agreement with the work of Crawford and Suchard [2013] whose simulations indicate that their approximate maximum likelihood procedure yields an unbiased consistent estimator of σ^2^. This is illustrated using the example of the Carnivora order studied previously by Crawford and Suchard [2013].

The phenotype modelled by a Brownian motion is usually interpreted as the case of neutral evolution with random oscillations around the ancestral state. This model was later developed into an adaptive evolutionary model based on the Ornstein–Uhlenbeck process by Felsenstein [1988], Hansen [1997], Butler and King [2004], Hansen et al. [2008], Bartoszek et al. [2012]. The tree-free setting using the Ornstein-Uhlenbeck process was addressed by Bartoszek and Sagitov [2012] where for the Yule-Ornstein-Uhlenbeck model, some phylogenetic confidence intervals for the optimal trait value were obtained via three limit theorems for the phylogenetic sample mean. Furthermore, it was shown that the phylogenetic sample variance is an unbiased consistent estimator of the stationary variance of the process.

At the end of their discussion Crawford and Suchard [2013] write that as the the tree of life is refined interest in “tree-free” estimation methods may diminish. They however indicate that “tree-free” estimates may be useful to calculate starting points for simulation analysis. We certainly agree with the second statement but believe that development of “tree-free” methods should proceed alongside that of “tree-based” ones.

One of the most useful features of the tree-free comparative models is that they offer a natural method of tree growth allowing for study of theoretical properties of phylogenetic models as demonstrated in this work [and also Sagitov and Bartoszek, 2012, Bartoszek and Sagitov, 2012, Bartoszek, 2014, Crawford and Suchard, 2013]. Another alternative to studying properties of these estimators is the tree growth model proposed by Ané [2008], Ho and Ané [2013], Ané et al. [2014]. In this setup the total height of the tree is kept fixed and new tips are added to randomly chosen branches. These two approaches seem to be in agreement, at least up to the second moments, since e.g. they agree on the lack of consistency of estimating *X*_0_. In Sagitov and Bartoszek [2012] we showed that under the Yule Brownian motion model Var 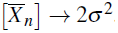.

In a practical situation “tree-free” methods can be used for a number of purposes. Firstly as pointed out by Crawford and Suchard [2013] they can be useful for calculating starting points for further numerical estimation procedures or defining prior distributions in a Bayesian setting. Secondly they have to be used in a situation where the tree is actually unknown e.g. when we are studying fossil data or trying to make predictive statements about future phenotypes, e.g. development of viruses. Thirdly they can be used for various sanity checks. If they contradict “tree-based” results this could indicate that the numerical method fell into a local maximum.

The paper has the following structure. Section 2 presents the model, the main results and an application. Section 3 states two lemmata and a proposition directly yielding the assertion of Theorem 2.1. Proposition 3.1 deals with the covariances between coalescent times for randomly chosen pairs of tips from a random Yule *n*-tree. The properties of the coalescent of a single random pair were studied previously by e.g. Steel and McKenzie [2001] and Sagitov and Bartoszek [2012]. In Section 4 we state two lemmata needed for the proof of Proposition 3.1. Section 5 contains two further lemmata and the proof of Proposition 3.1. In Section 6, 7, and 8 we prove the lemmata from Sections 3, 4, and 5. Appendix A contains some useful results concerning harmonic numbers of the first and second order.

## 2 The main results

The basic evolutionary model considered in this paper is characterized by four parameters (*λ, n,X*_0_, σ^2^) and consists of two stochastic components: a random phylogenetic tree defined by parameters (*λ, n*) and a trait evolution process along a lineage defined by parameters (*X*_0_, σ^2^). The first component, species tree connecting n extant species, is modelled by the pure birth Yule process [Yule, 1924] with the birth (speciation) rate λ and conditioned on having *n* tips [Gernhard, 2008]. For the second component we adapt the approach by assuming that for a given *i* = 1,…, *n*, the current trait value 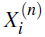 has evolved from the ancestral state *X*_0_ according to the Brownian motion with the local variance σ^2^.

Treating the collection of the current trait values 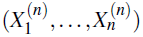 generated by such a process as a sample of identically distributed, but dependent, observations, we are interested in the properties of the basic summary statistics

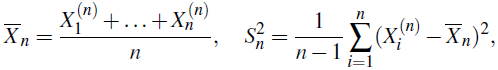

the sample mean and sample variance.

According to [Sagitov and Bartoszek, 2012] we have

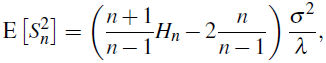

see Fig 1, left panel (all simulations are produced using the TreeSim [Stadler, 2009, 2011] and mvSLOUCH [Bartoszek et al., 2012] R packages). It follows that the normalized sample variance

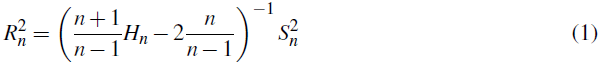

gives an unbiased estimator of the compound parameter 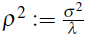 for the Yule-Brownian-Motion model, see Fig 2. In the comparative phylogenetics framework the ratio *ρ*^2^ can be called the *evolutionary rate* as it measures the speed of change in the trait value when the time scale is such that we expect one speciation event per unit of time and per species. The next theorem is the main asymptotic result of this paper, illustrated by Fig 1, right panel.

**Figure 1:**
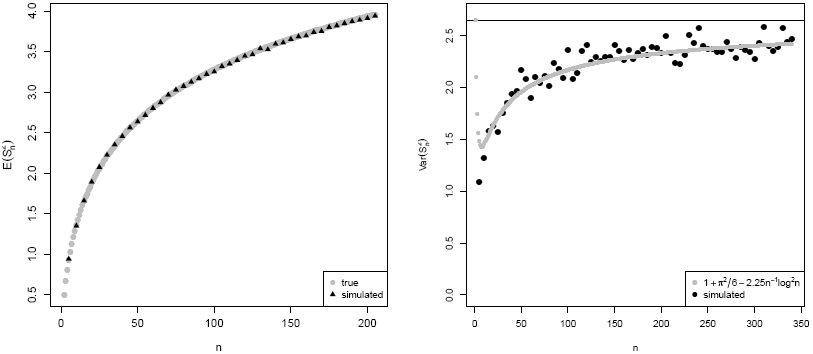
Left: True and simulated values of E [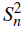], right: simulated values of Var [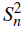] with limit equalling π^2^/6+ 1. Each point comes from 10000 simulated Yule trees and Brownian motions on top of them. Parameters used in simulations are *λ =* 1, *X*_0_ *=* 0 and σ^2^= 1. The grey line on the right panel fits a curve based on the convergence rate O (*n^−1^* log *n*^2^).

**Figure 2:**
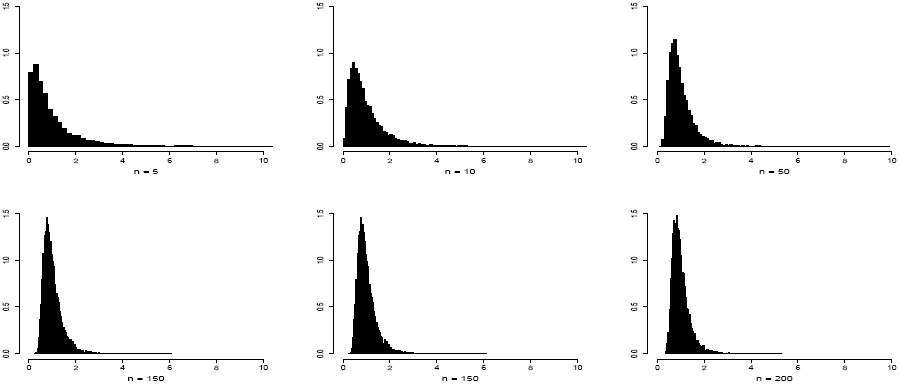
Histograms of 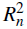 for left to right top *n =* 5,10,50 and bottom *n =* 100,150,200. Parameters used in simulations are *λ =* 1, *X*_0_ *=* 0 and σ^2^ = 1.

### Theorem 2.1

*Consider the sample variance 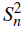 for the Yule-Brownian-Motion model with parameters (λ,n,X_0_,σ^2^). Its variance satisfies the following asymptotic relation*

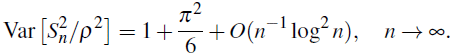

In terms of our estimator (1), Theorem 2.1 yields

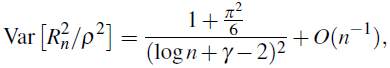

where *γ* = 0.577 is the Euler constant, implying that 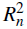 is a consistent estimator of the evolutionary rate *ρ*^2^. It follows that for large n, the standard error (estimated standard deviation) of the unbiased estimator 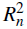 can be approximated by

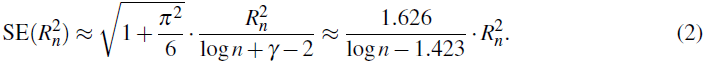

The estimator of Eq. (1) should be compared to the approximate maximum–likelihood estimator for the local variance σ^2^ recently proposed by Crawford and Suchard [2013] in the same framework of the Yule–Brownian–Motion model. The main difference between two approaches is that in Crawford and Suchard [2013] it is assumed that one knows both the number of tips and the total height of the otherwise unknown species tree. The Crawford-Suchard estimator is based on a closed form of the distribution of phylogenetic diversity – the sum of branch lengths connecting the species in a clade.

As an application of their estimator, Crawford and Suchard [2013] study different families of the Carnivora order, estimating σ^2^ for each of the 12 clades. The data for the log-body-size disparities was taken from the PanTHERIA database [Jones et al., 2009]. The data summary and the Crawford-Suchard estimates are shown in the left part of Tab. 1. In the right part of Tab. 1 we present our estimates 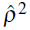 for the evolutionary rate parameter *ρ*^2^ = σ^2^/λ for each of the 12 families in the Carnivora order. The standard error is computed using (2). We note that the data does not take into account the newly described species *Bassaricyon neblina* from the Procyonidae family [Helgen et al., 2013].

In the next-to-last column we list the ratios demonstrating a surprisingly good agreement between our and Crawford-Suchard estimates. The ratio is taken between two products: 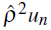 on one hand, and 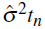 on the other. Here *u_n_ =* E [*U_n_*] is the expected age of the conditioned standard Yule process with *λ =* 1, while *t_n_* is the clade age assumed to be known in the Crawford-Suchard framework. Both 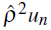 and 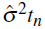 estimate the same quantity - the variance in the trait values for the evolution of the corresponding clade. Therefore, one should expect these ratios to be close to one. And indeed, the 12 ratios have mean 0.97 and standard deviation 0.20.

Our estimator and its standard error are computed by simple formulae given above. A major weakness of our estimator is relatively big standard error for realistic richness values, see the 7th column in Tab. 1. This can be explained by the fact that we do not use an additional information about the species tree, like the height of the tree used in the Crawford-Suchard estimator.

This close agreement is obtained despite a number of features that complicates the comparison between two methods. Our approach in its current form does not allow to take into account the fact that some trait values are missing. We calculated 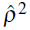 for the trait disparity as if it was computed using all *n* trait values. Moreover, it is not be clear how to take into account the measurement variance. As shown by Hansen and Bartoszek [2012] even with a known tree, the measurement error can cause very diverse effects. Therefore we would expect the situation to be even more interesting when we integrate the phylogeny out.

**Table 1:** Data summary. 2nd column: clade richness (number of missing trait values); 3rd column: the clade age in millions of years; 4th column: 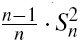 trait disparity; 6th column: the expected age *u_n_ =* E [*U_n_*] of the conditioned standard Yule process with *λ =* 1.

| Family | $n$ | $t_n$ | Disparity | $\hat{\sigma}^2$ (SE) | $u_n$ | $\hat{\rho}^2$ (SE) | $\frac{\hat{\rho}^2 u_n}{\hat{\sigma}^2 t_n}$ | $\frac{\hat{\rho}^2}{\hat{\sigma}^2 / \lambda}$ |
| --- | --- | --- | --- | --- | --- | --- | --- | --- |
| Felidae | 40 (7) | 33.3 | 1.588 | .080 (.009) | 4.279 | .649 (.466) | 1.042 | 0.560 |
| Viverridae | 35 (6) | 37.4 | 0.662 | .029 (.004) | 4.147 | .284 (.217) | 1.086 | 0.676 |
| Herpestidae | 33 (4) | 25.5 | 0.482 | .030 (.003) | 4.089 | .211 (.166) | 1.128 | 0.485 |
| Eupleridae | 8 (0) | 25.5 | 0.916 | .079 (.010) | 2.718 | .758 (1.72) | 1.023 | 0.662 |
| Hyaenidae | 4 (0) | 32.2 | 0.805 | .122 (.005) | 2.083 | .999 (19.5) | 0.530 | 0.565 |
| Canidae | 35 (3) | 48.9 | 0.678 | .030 (.004) | 4.147 | .290 (.221) | 0.825 | 0.667 |
| Ursidae | 8 (0) | 42.6 | 0.303 | .024 (.002) | 2.718 | .251 (.569) | 0.667 | 0.722 |
| Otariidae | 16 (2) | 24.5 | 0.386 | .028 (.003) | 3.381 | .227 (.274) | 1.119 | 0.559 |
| Phocidae | 19 (0) | 24.5 | 0.751 | .052 (.005) | 3.548 | .410 (.438) | 1.142 | 0.544 |
| Mephitidae | 12 (3) | 32.0 | 0.570 | .039 (.005) | 3.103 | .384 (.588) | 0.955 | 0.679 |
| Mustelidae | 59 (10) | 27.4 | 2.263 | .126 (.014) | 4.663 | .811 (.497) | 1.095 | 0.444 |
| Procyonidae | 14 (1) | 27.4 | 0.531 | .037 (.004) | 3.252 | .332 (.444) | 1.065 | 0.619 |

In their work Crawford and Suchard [2013] estimated the overall speciation rate to be 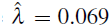 per million years. The last column of Tab. 1 demonstrates that using this common value for the speciation rate *λ* produces huge discrepancy between our estimates 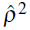 for the rates of evolution *ρ*^2^ = σ^2^/λ and the rates of evolution computed using the Crawford-Suchard estimates for σ^2^. This observation points out that a fair direct comparison of 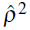 and 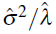 would requires specific estimates of the speciation rate *λ* for each of the 12 clades.

## 3 Outline of the proof of Theorem 2.1

We start with a general observation, Lemma 3.1, concerning the sample variance

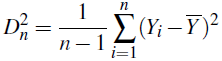

of *n*, possibly dependent and not necessarily identically distributed, observations (*Y*_1_,…, *Y*_n_) with sample mean 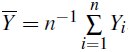

### Lemma 3.1

*If* (*W*_1_, *W*_2_, *W*_3_, *W*_4_) *is a random sample without replacement from random values* (*Y*_1_,…, *Y*_*n*_), *then*

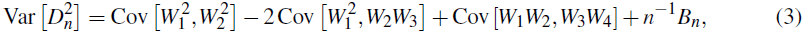

where

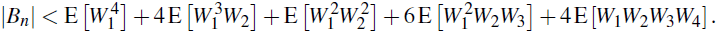

Observe that in terms of the sample variance for the scaled trait values

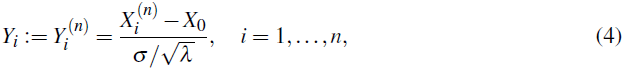

we have 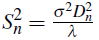, and to prove Theorem 2.1 we have to verify that

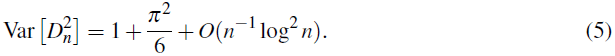

The Yule *n*-tree underlying the set of scaled values (4) has unit speciation rate. We call it the standard Yule *n*-tree, and denote by 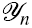 be the σ–algebra generated by all the information describing this random tree. Under the Brownian motion assumption the trait values (4) are conditionally normal with

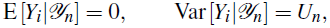

where *U*_n_ is the height of the standard Yule *n*-tree, see Fig. 3. Moreover, see Section 6, we have

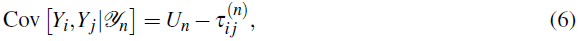

where 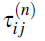 is the backward time to the most recent common ancestor for a pair of distinct tips (*i, j*) in the standard Yule *n*-tree, see Fig. 3. For a quadruplet (*i, j, k, l*) of tips randomly sampled without replacement out of *n* tips in the standard Yule *n*-tree, we denote

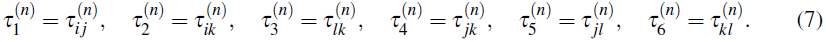

**Figure 3:**
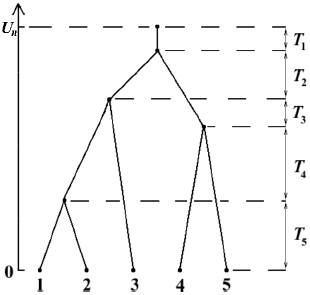
An example of a standard Yule *n*-tree with *n* = 5. The tree height is *U*_*n*_ = *T*_1_ +…+ *T*_*n*_, where *T*_*i*_ are the times between the consecutive speciation events. The 10 pairwise coalescent times 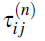 for the tips of the tree are 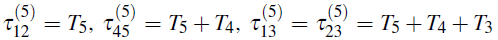, and 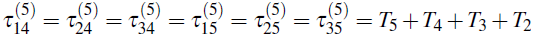.

### Lemma 3.2

*Let* (*W*_1_, *W*_2_, *W*_3_, *W*_4_) *be a random sample without replacement of four trait values out of n random values defined by* (4). *Then in terms of the coalescent times* (7) *we have*

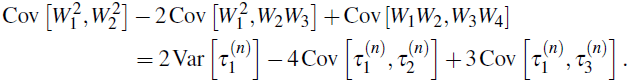

In view of Lemmata 3.1 and 3.2 which are proven in Section 6, to verify (5) it suffices to show the following asymptotic result.

### Proposition 3.1

*Consider the coalescent times* (7). *As n* → ∞,

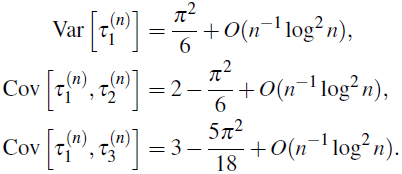

Notice that the key Proposition 3.1 concerns only the first component of the evolutionary model we study - the standard Yule *n*-tree. For the standard Yule *n*-tree it is well known that the times between the consecutive speciation events (*T*_1_,…, *T*_*n*_) are independent exponentials with parameters (1,…, *n*) respectively, see Fig. 3. As shown in Gernhard [2008], this property corresponds to the unit rate Yule process conditioned on having *n* tips at the moment of observation, assuming that the time to the origin has the improper uniform prior [see also Feller, 1971].

## 4 Coalescent indices of the standard Yule *n*-tree

Following the standard Yule *n*-tree from its root toward the tips we label the consecutive splittings by indices 1,…, *n*-1: splitting *k* is the vertex when *k*-1 branches turn into *k* branches. We define three random splitting indices (as we interested in four randomly chosen tips out of *n* available):

- *K*_*n*_ is the index of the splitting where two randomly chosen tips coalesce,
- *L*_*n*_ be the index of the splitting where the first coalescent among three randomly chosen tips takes place,
- *M*_*n*_ be the index of the splitting where the first coalescent among four randomly chosen tips takes place.

To avoid multilevel indices in the forthcoming formulae, we will often use the following notational convention

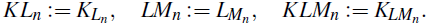

To illustrate these indices, turn to the Fig. 3. If the two randomly chosen tips are (1,2), then *K*_*n*_ = 4. If the three randomly chosen tips are (2,3,4), then *L*_*n*_ = 4, *K*_*L*_*n*__ = 2. If the four randomly chosen tips are (2,3,4,5), then *M*_*n*_ = 3, *L*_*M*_*n*__ = 2, *K*_*LM*_*n*__ = 1.

The importance of these random indices comes from the following representations. Denote 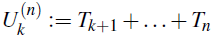 the sum of adjacent times between splittings in the Yule tree. Clearly,

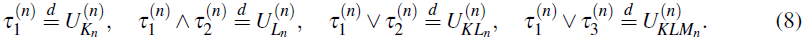

To prove Proposition 3.1 we need to know the distributions of these random splitting indices. The next two lemmata giving these distributions are proved in Section 7.

### Lemma 4.1

*Then*

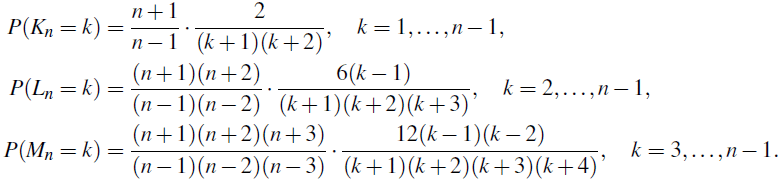

### Lemma 4.2

*The random numbers K*_*L*_*n*__, *L*_*M*_*n*__, *K*_*LM*_*n*__ *have the following distributions*

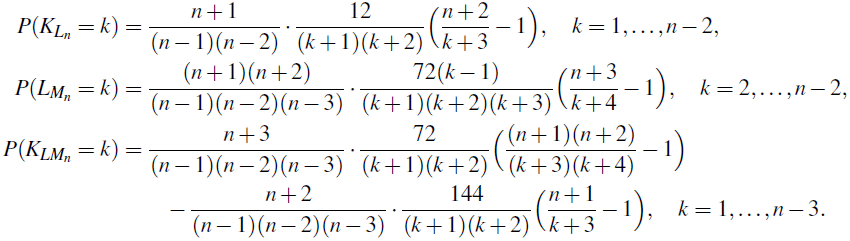

## 5 Proof of Proposition 3.1

In view of (8), the harmonic numbers

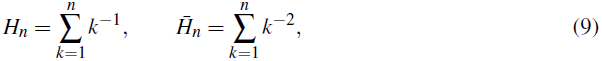

play an important role in our calculations as

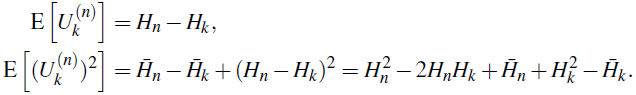

### Lemma 5.1

*We have*

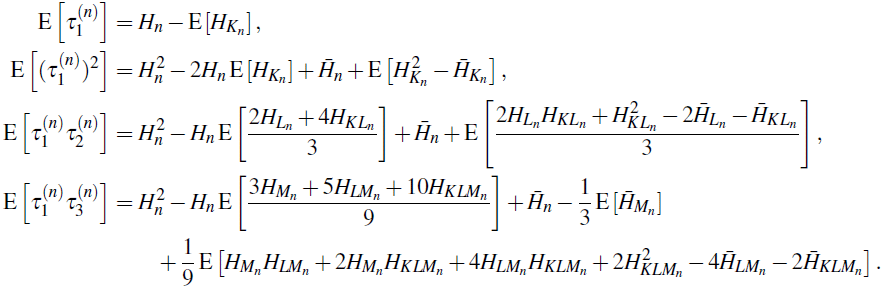

In view of Lemma 5.1, proven in Section 7, the asymptotic results stated in Proposition 3.1 are computed using the following relations involving the harmonic numbers (9).

### Lemma 5.2

*We have as n* → ∞

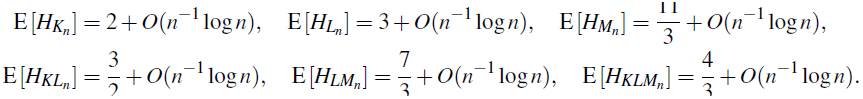

### Lemma 5.3

*Let a*_*n*_ ⇉ *a stand for a*_*n*_ = *a*+O(*n*^−1^ log^2^ *n*) *as n* → ∞. *Then*

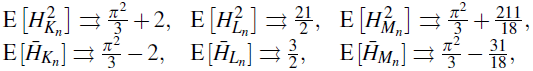

*and*

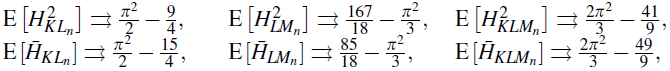

*and*

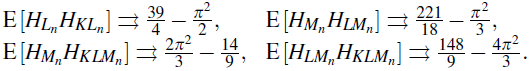

The proofs of the last two lemmata are given in Section 8 using the auxiliary results from Appendix A.

With Lemmata 5.1 - 5.3 at hand, the remaining proof of Proposition 3.1 is straightforward. The first statement

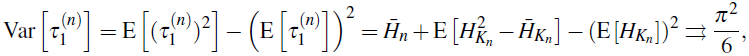

is obtained applying the classical relation 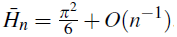. Further, Lemma 5.1 yields

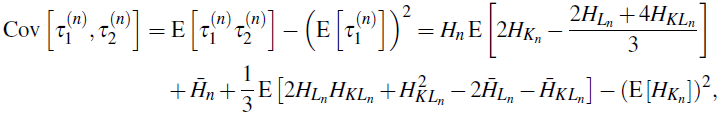

where according to Lemma 5.2

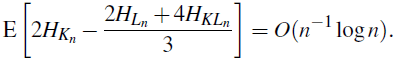

Thus, applying Lemma 5.3 we obtain the second statement

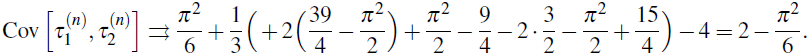

Finally, the third statement follows from

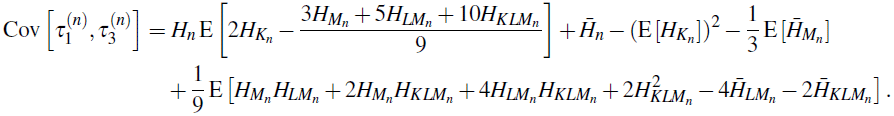

Indeed, according to Lemma 5.2

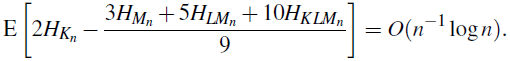

Moreover, from the following three limits

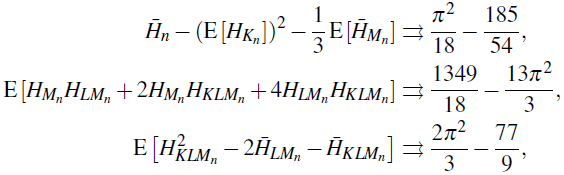

we get the stated overall limit

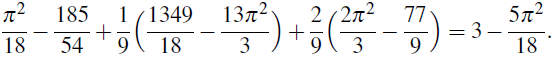

## 6 Proofs of Lemmata 3.1 - 3.2

### Proof of Lemma 3.1.

Using the representation

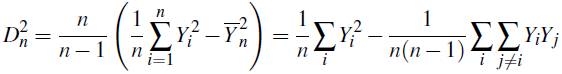

we find that

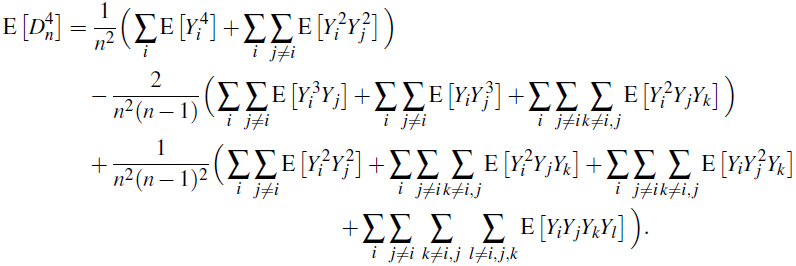

If (*W*_1_, *W*_2_, *W*_3_, *W*_4_) is a random sample without replacement of four out of *n* trait values, then

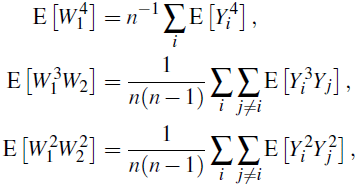

and

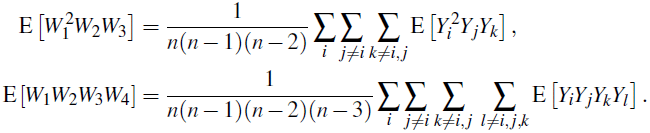

Therefore, we have

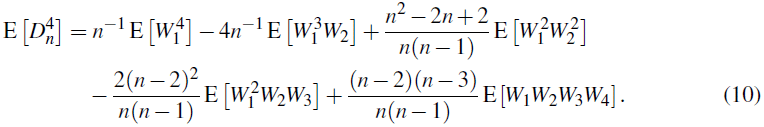

Since

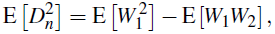

we conclude

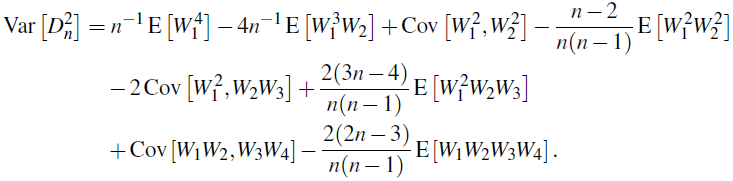

The stated relations follow with

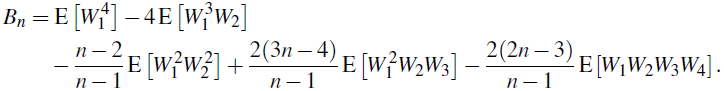

### Proof of Lemma 3.2.

Denote by 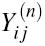 the normalized trait value of the most recent common ancestor of the tips (*i, j*). Let 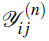 stand for the σ–algebra generated by the pair 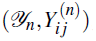, then

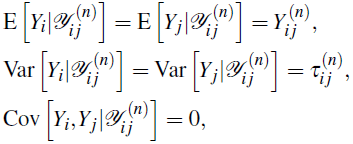

implying (6)

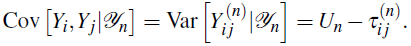

By Eq. (13) of Bohrnstedt and Goldberger [1969], we have

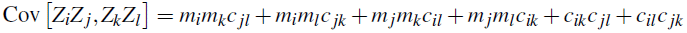

for any sequence of normally distributed random values Z_1_,Z_2_,… with means E[Z_i_] = *mi* and covariances Cov [*Z_i_,Z_j_*] = *c_ij_*. In the special case with *mi* **=** 0 it follows

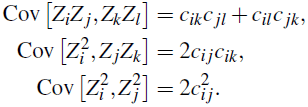

Using conditional normality of *Y_i_* and putting *c*_*ij*_ = *U*_*n*_ -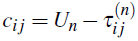, we derive from these relations that

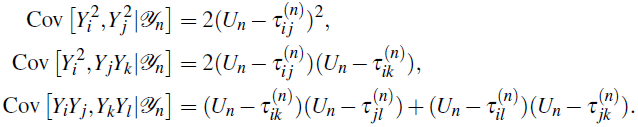

yielding in terms of (7),

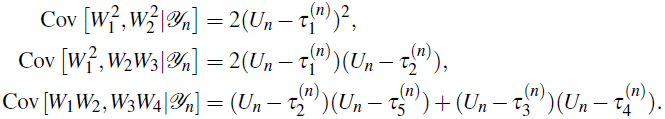

By the total covariance formula, we derive

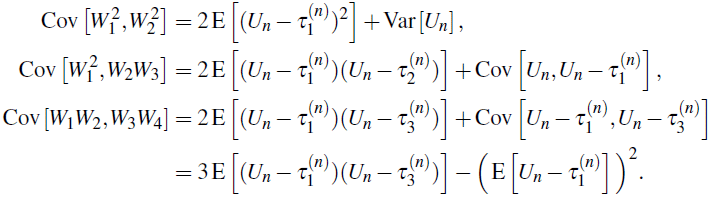

Combining these relations we get

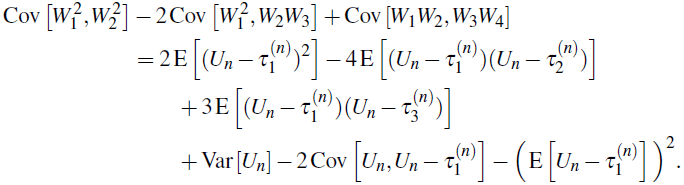

This together with

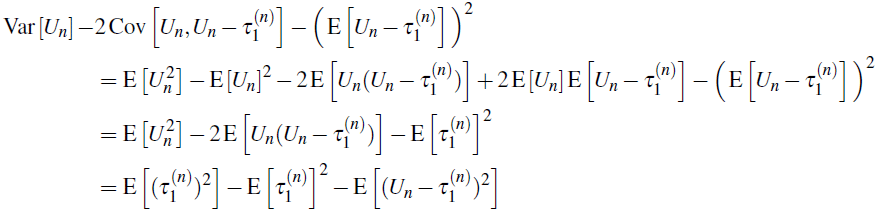

implies the assertion of the Lemma 3.2

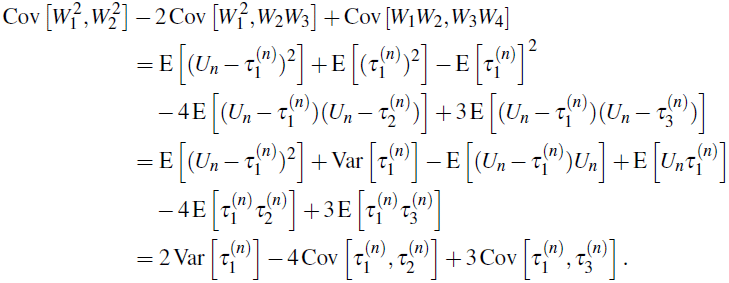

## 7 Proofs of Lemmata 4.1, 4.2, and 5.1

### Proof of Lemma 4.1

From the definition of *K_n_* it is easy to see that, for *k = 2,…, n,*

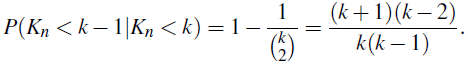

Therefore,

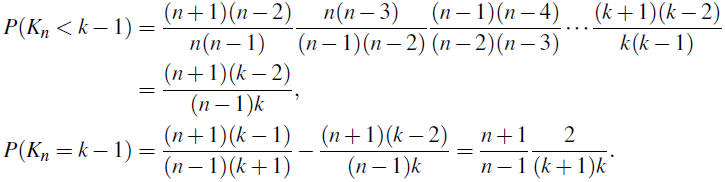

Similarly, for *k* = 3,…, *n*,

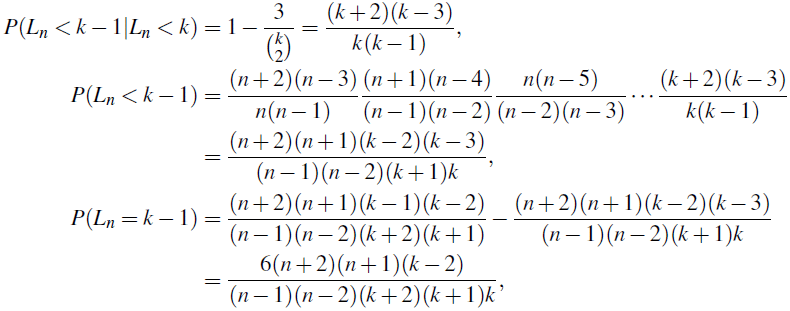

and, for *k* = 4,…, *n*,

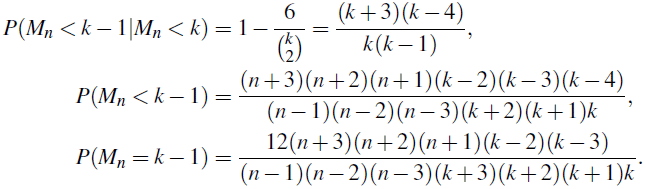

### Proof of Lemma 4.2

Clearly,

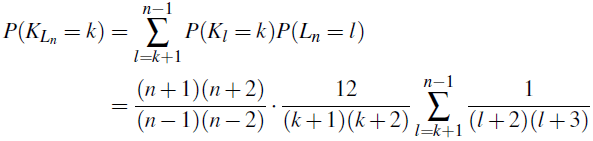

leads to the first assertion. Further,

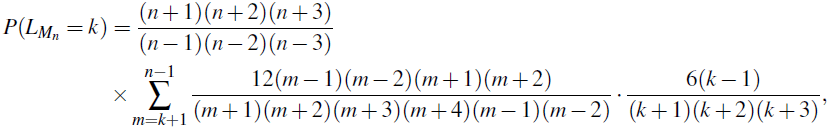

and

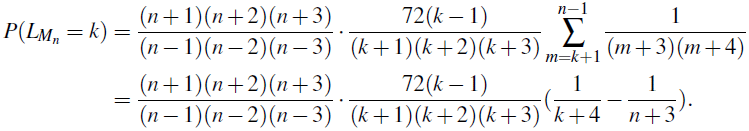

Finally,

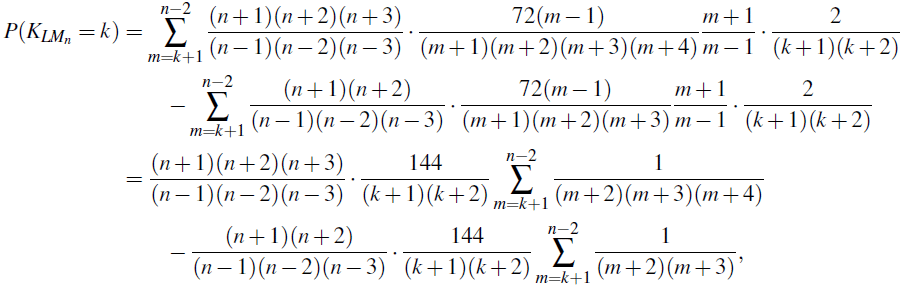

and therefore, it remains to use the equalities

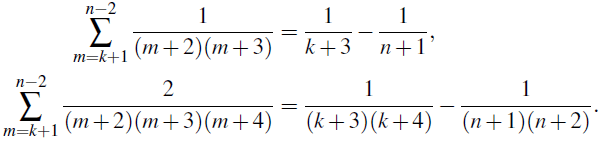

### Proof of Lemma 5.1.

We have

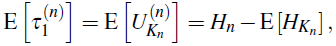

and

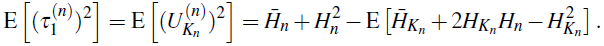

Further, using (8) and 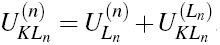, we get

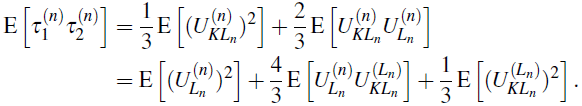

Thus

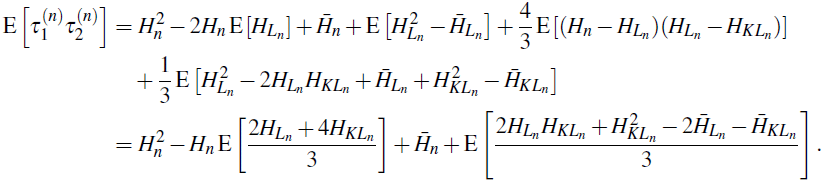

Finally, for two pairs of sampled tips, we have three coalescent events to consider: going from four to three selected nodes, 4 → 3, going from three to two selected nodes, 3 → 2, and going from two to one selected nodes, 2 → 1. The coalescent 4 → 3 holds across the two pairs with probability 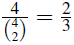 and within a pair with probability 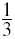. Given the former outcome, the coalescent 3 → 2 holds again across the pairs with probability 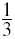 and within a pair with probability 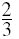. Otherwise, the coalescent 3 → 2 holds across the pairs with probability 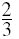 and within the second pair with probability 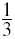. The four possibilities 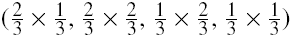 produce the following four terms in

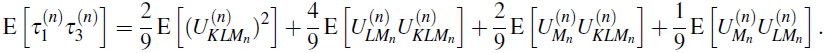

It follows,

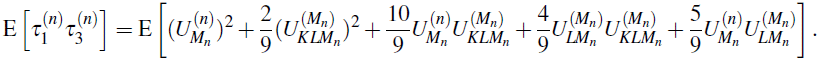

Using the representation for 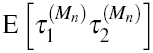

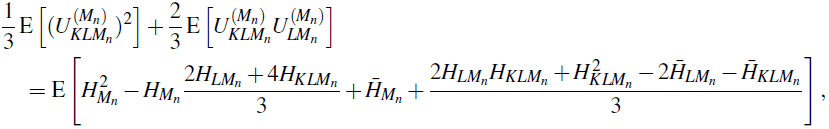

we can write

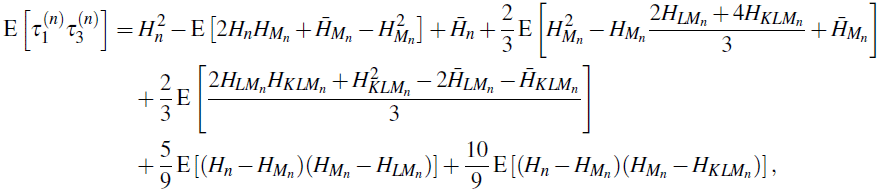

which after a rearrangement gives the last statement.

## 8 Proof of Lemmata 5.2 - 5.3

In this section we will often use the elementary relations of the following type

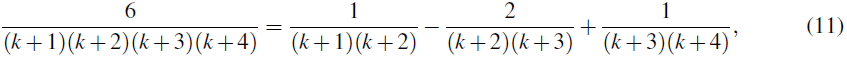

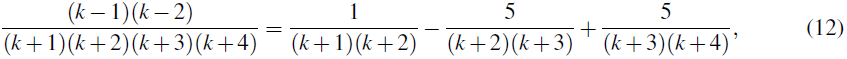

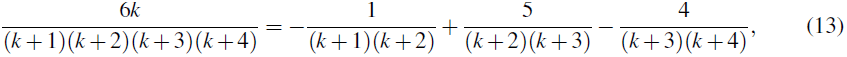

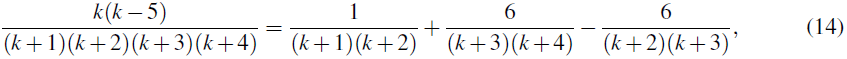

valid for all *k* ≥ 1.

### Proof of Lemma 5.2.

The first three stated relations are obtained using Lemmata 4.1 and A.1.

Equalities

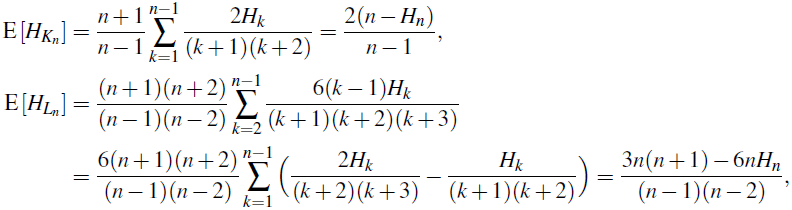

give the first and the second stated relations, and the third one follows from

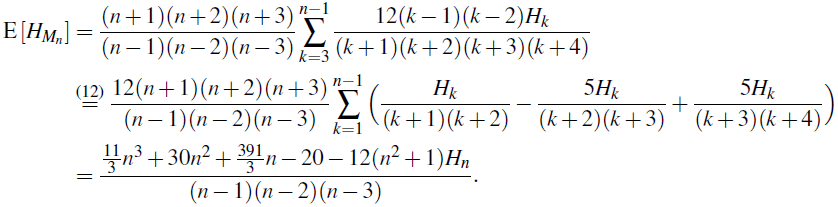

The second three stated relations are obtained similarly using Lemmata 4.2 and A.1. Indeed,

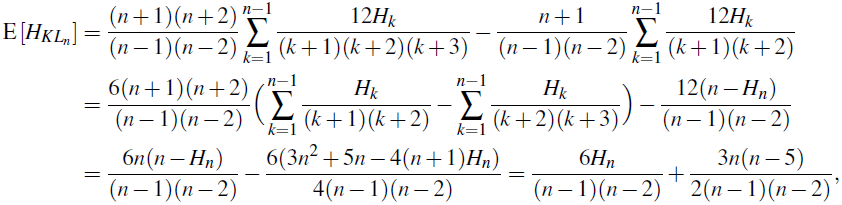

implying 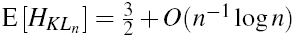. Furthermore,

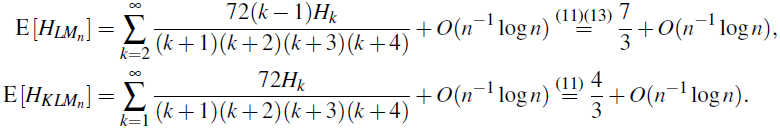

### Proof of Lemma 5.3.

The stated relations are obtained using Lemmas 4.1, 4.2, A.1, A.2. Firstly,

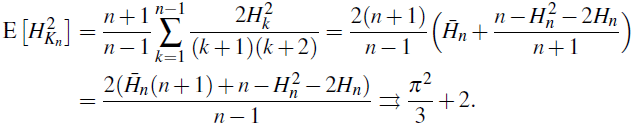

Similarly, we have

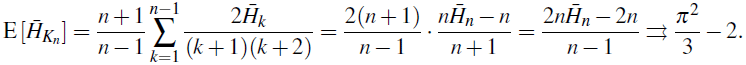

Observe that the limit is 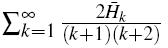. In the same manner we obtain

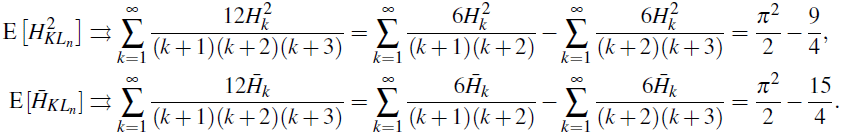

Using the decomposition (12) we find

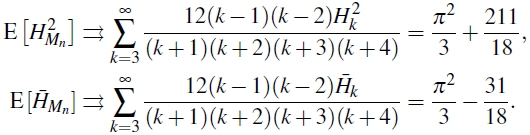

Using the difference between (13) and (11) we find

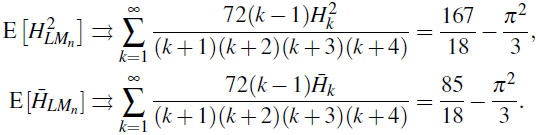

Using (11) we find

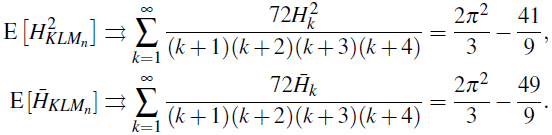

Since 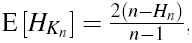, we have

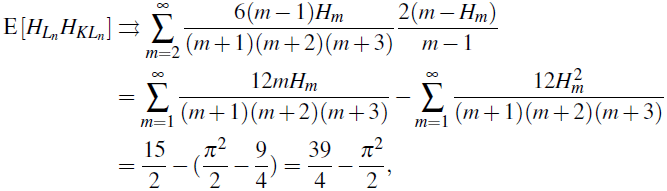

where we use the following corollary of Lemma A.1

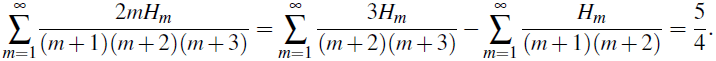

Similarly,

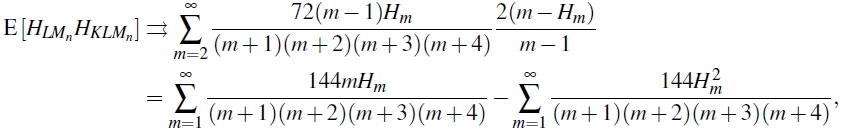

where

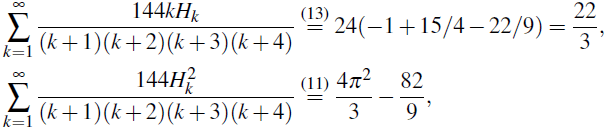

so that 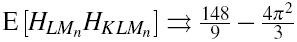.

Further, in view of 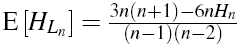, the limit for E[*H*_*M*_*n*__*H*_*LM*_*n*__] can be computed as

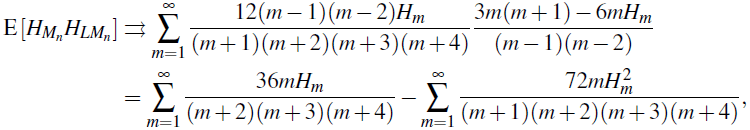

where

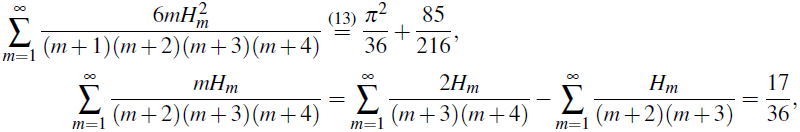

yielding 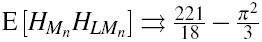. Finally, from

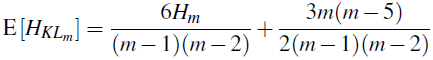

we get

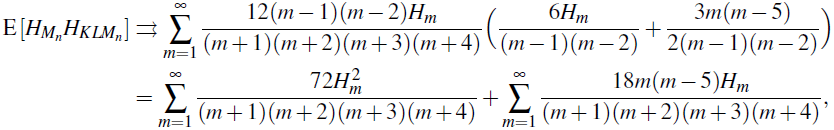

where

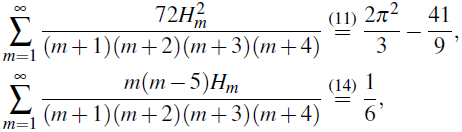

so that 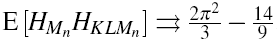.

## Acknowledgements

The research of Serik Sagitov was supported by the Swedish Research Council grant 621-2010-5623. Krzysztof Bartoszek was supported by the Centre for Theoretical Biology at the University of Gothenburg, Svenska Institutets Östersjösamarbete scholarship nr. 11142/2013, Stiftelsen för Vetenskaplig Forskning och Utbildning i Matematik (Foundation for Scientific Research and Education in Mathematics), Knut and Alice Wallenbergs travel fund, Paul and Marie Berghaus fund, the Royal Swedish Academy of Sciences, and Wilhelm and Martina Lundgrens research fund.

## A Auxiliary results involving harmonic numbers

Some of the following results can be found in Adamchik [1997] and Sofo [2011, 2012, 2013].

### Lemma A.1

*We have*

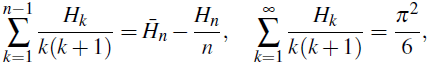

*and for m* ≥ 1,

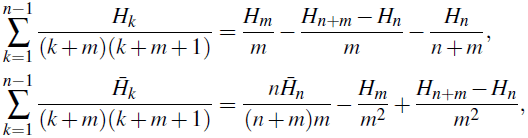

*so that*

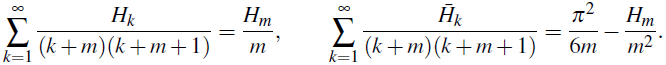

*In particular,*

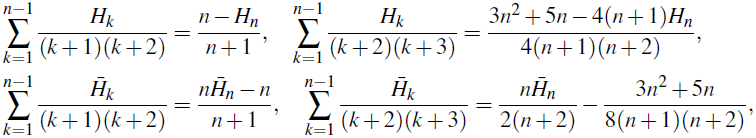

*and*

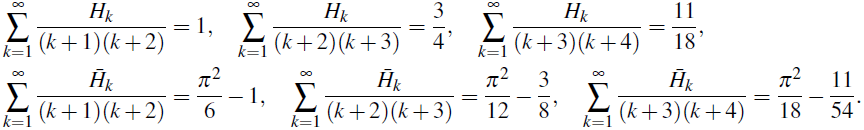

PROOF Clearly,

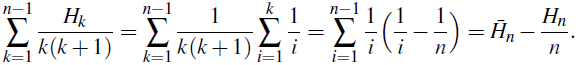

Similarly for *m* ≥ 1, we have

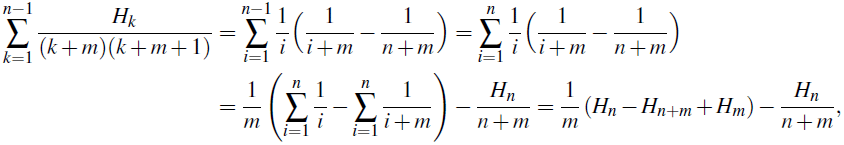

and

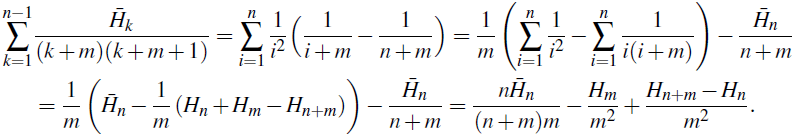

### Lemma A.2

*We have*

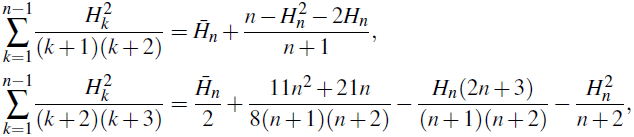

*and generally for m* ≥ 1,

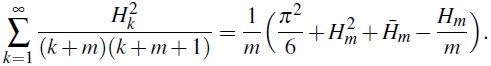

*In particular,*

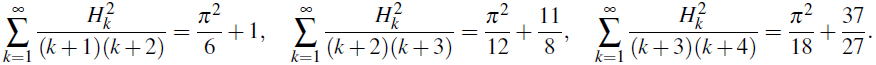

PROOF For *m* ≥ 1,

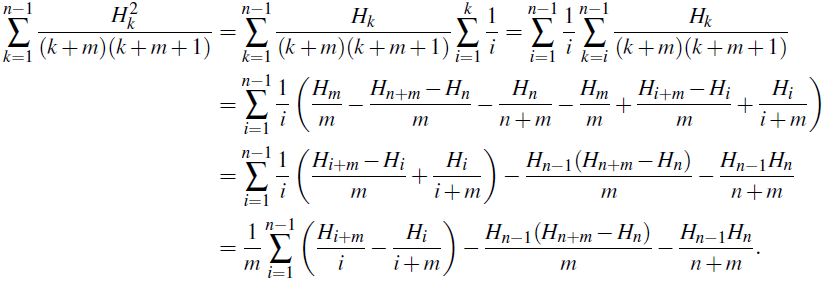

Observe that

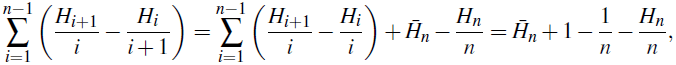

and for *k* ≥ 2,

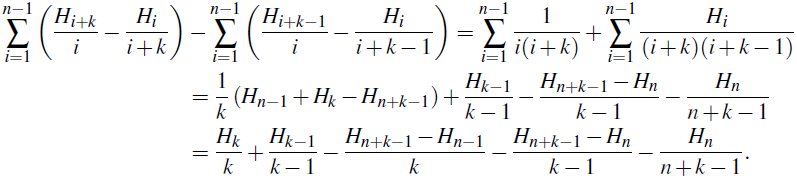

It follows

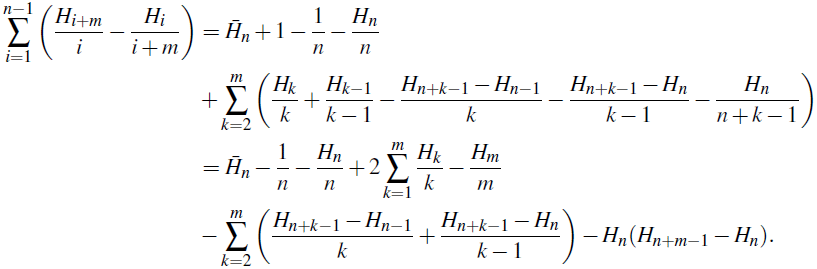

Using the classical relation 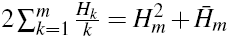 which follows from

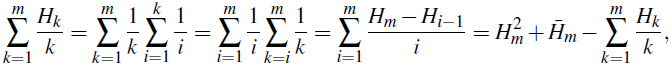

we get

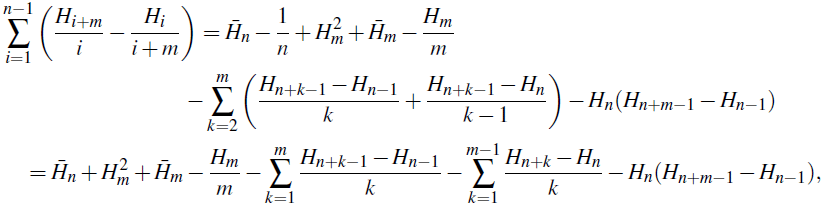

Thus

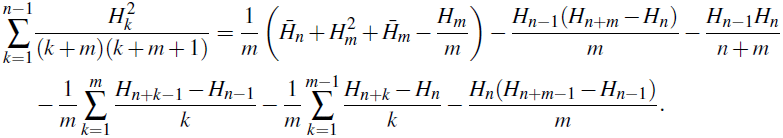

To finish the proof it remains to observe that 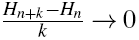 as n → ∞ for any fixed *k*.

